# Live Mouse Tracker MCP: Custom behavioral analysis accessible to all

**DOI:** 10.64898/2026.11.20.768067

**Authors:** Benoît Forget, Guillaume Nœtinger, Xavier Mousset, Manon Moreau, Mathieu Matsima, Nathalie Lemière, Sabrina Coqueran, Thomas Bourgeron, Fabrice de Chaumont

## Abstract

We previously developed the Live Mouse Tracker (LMT) to explore the wide repertoire of social interactions in wild-type mice and in models of autism. This tool records the positions and annotates the behaviors of groups of mice over days to months for quantitative behavioral science, but analysing these rich data usually requires custom code. As a result, less than 10% of LMT users analyse their data autonomously. To make data analysis accessible to every user, we present LMT-MCP, which includes a Model Context Protocol (MCP) server that connects any local LMT database to consumer large language models (LLMs). Users can now run their analyses by asking questions in natural language, without writing any code. To our knowledge, this is the first use of an MCP server in behavioral science. We illustrate the approach with two scenarios: an AI-assisted data exploration and a comparative study of homozygous and heterozygous *Shank3* mutant mice. The results are compared with existing reports produced without AI to demonstrate the robustness of the method and its results. Guidelines and system prompts are integrated to ensure these analyses remain valid and reproducible.

## Introduction

Quantitative behavioral studies using murine models are essential for investigating social phenomena, for instance to gain a deeper understanding of addiction [1], social hierarchies [2], or psychiatric conditions [3,4] in a social context. Such experiments typically involve tracking and labelling the behavior of a group of mice during tests lasting from a few hours to several days or months. To address this need, we created the [Live Mouse Tracker](https://micecraft.org/lmt/) (LMT) [5] and a dedicated lmt-analysis package to process data. Then, the community created generic graphical frontend tools to help analyze the large amounts of data produced. Tools such as LMT-Eye, [LMT toolkit analysis](https://github.com/ntorquet/lmt_toolkit_analysis), LMT-easy, [LMT Widget tools](https://pypi.org/project/LWTools/), [Tripping rats](https://github.com/terelacco/tripping_rats) or [MouseKing](https://github.com/DaleAnnear/MouseKing) provided different angles of analysis of the data. Yet, given the richness of the LMT data, which contain the tracking of animals, the labelling of the behaviors plus additional TTL inputs or synchronisation with other devices or programs, a growing number of users wish to request new types of analysis. Each time, this requires custom development of an analysis platform. Consequently, experimentalists must master a wide range of coding tools. However, the programming skills involved usually lie outside the typical training or background of the behavioral science community as it has been pointed out in other biological science communities [6]. Seven years after the release of the LMT, a survey of our registered users shows that less than 10% can perform their data analysis autonomously or have access to a data analyst.

To bridge the gap that has emerged in biology between the design and execution of experiments and the scientists expected to use and interpret them, artificial intelligence (AI) tools have already been proposed [7] and listed [8]. For instance, Large Language Models (LLMs) have been crafted for automatic labelling of videos [9,10]. Here, we present a comprehensive experimental pipeline that culminates in a Model Context Protocol (MCP) [11], a standard developed to interface any AI model to a local database. This ready-made solution brings the power of consumer LLMs to LMT users who can now prompt their data analysis in natural language within minutes. MCPs have been deployed successfully in other biological domains such as spatial transcriptomic [6], neuroimaging [12] or to make bibliography more interactive [13] but, to our knowledge, this is a first in behavioral science. This leads to a considerable saving of time, engineering skills and effort. To illustrate its use and limitations, we provide two use case scenarios: one in which we perform data exploration with LLM, and a second one in which two datasets of *Shank3* heterozygous and homozygous mutant mice are compared to wild type mice. The results obtained using the LMT-MCP are then compared with our previous analysis to assess the robustness of the method.

While LMT-MCP provides a shortcut for behavioral scientists, its flexibility also makes it easy to run unstructured or non-reproducible analyses [14,15,16]. We therefore propose a set of guidelines to standardize its use, along with pre-prompts to ensure that statistics are generated, and scripts are available for transparency and reproducibility. Far from replacing the work of research engineers, it allows them to focus on more specific needs or, on the contrary, more general considerations to expand the scope of the LMT platform.

## Results

### MCP, software and installation tutorial

Here we provide information on what is an MCP application, the software architecture and how to download and install the software. Direct access to online how-to here: https://micecraft.org/lmt/mcp/

### The MCP Application

The Model Context Protocol (MCP) is an open standard that lets an LLM assistant connect to external software and data. An MCP server exposes a set of tools **(Fig.1)**, such as reading an animal list or counting events, which the language model can call while it answers a question. In our implementation, the MCP is local to the user computer. The LLM can write scripts that are run locally, by the user’s computer. Our LMT-MCP server is distributed as a standalone executable for Windows, so it can be installed without any Python environment. At the start of each session, the server also sends the language model a short text of `INSTRUCTIONS`, which explains to the LLM in natural language the conventions of LMT data. For the benefit of our users, we set these instructions to require the language model to return the Python analysis scripts it used (see supplementary materials: instructions). These scripts run offline on the database, so that any result can be checked and reproduced without the AI. We also state in the instructions that animals together in the same experiment/cage are not independent and that the experiment is the statistical unit, so that the model chooses appropriate statistical tests. We also depict how the whole data structure is made, explaining the field of the database and different pitfalls. By generating this instruction set, the LLM can justify what method to use to the user by exposing LMT guidelines. We will keep releasing new versions of the server that add tools, prompts and good-practice guidelines based on user feedback and needed support on discord: https://discord.gg/zWDHNf9eHM. A step by step installation tutorial is available here: https://micecraft.org/lmt/mcp/. The MCP free and under GPLv3 license, hosted on GitHub: https://github.com/gnoetinger/lmt-eye

**Figure 1.**
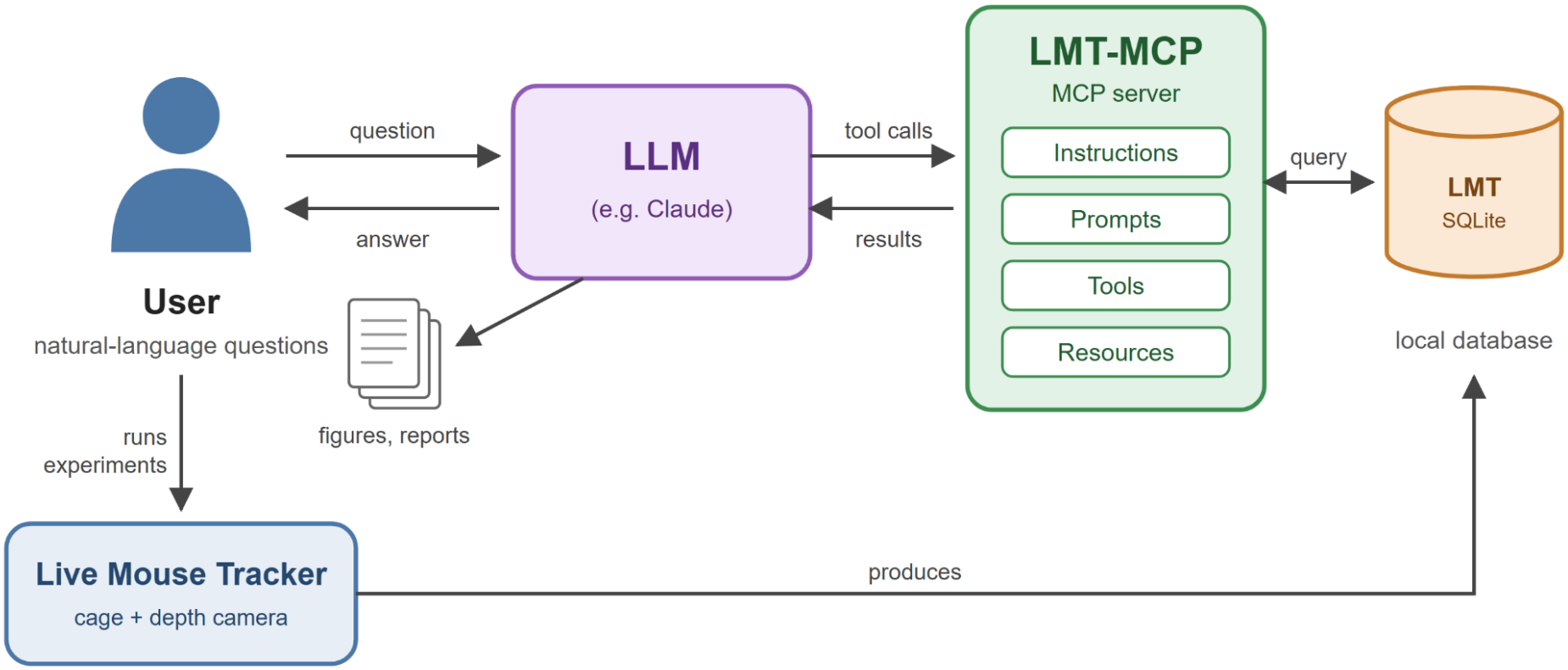
LMT-MCP architecture. The user runs experiments with the Live Mouse Tracker (LMT), which stores tracking and behavioral events in a SQLite database. The user asks questions in natural language to an LLM, which queries the database through the LMT-MCP server (instructions, prompts, tools, resources) and returns answers, figures and reports.

### Software architecture

LMT-Eye is a *Python* software designed to explore data from LMT. It loads a database and is able to generate reports displayed as an .*html* page which contains a variety of information on the activity of the mice. It is downloadable from [GitHub](https://github.com/gnoetinger/lmt-eye/tree/mcp-server) and extensively described in its [wiki](https://github.com/gnoetinger/lmt-eye/wiki) page. It was conceived to be customized for each specific application by manually coding a custom type of event.

Next to this preexisting program, we added the extension based on the MCP standard [11] that enables an AI to have access to a previously loaded database. This allows LLM to have access to the LMT database with the data staying local. Furthermore, it can access specific tools from LMT-Eye to access and analyse the database. Thus, contrary to just feeding the database to an LLM with no prior information, it benefits from the precise labelling and all the tools developed beforehand included in the LMT-Eye library. The MCP was added in this software to allow for a comparison by the user with the reports generated by the code custom-developed previously on LMT-Eye.

### Software installation

The program has been written in [Python](https://github.com/gnoetinger/lmt-eye.git). To ease its use, a Windows installer is provided. A tutorial for the installation is provided here: https://micecraft.org/lmt/mcp/.

### Use cases

In the following, we depict the use of LLM connected via MCP to LMT data. We used an LLM accessible to anyone: assistant Claude Code Opus 5.5 with a Pro plan on low effort. The LMT data is simply a file with the .sqlite extension, no server is needed to read it. The first scenario #1 is a “first contact with data” scenario, where one is discovering the data and is asking broad questions on a small database in an explorative way. It starts with simple questions getting gradually more and more complex up to the redaction of a report. The second scenario is based on a dataset of 39 experiments of 47h that has been previously analysed by our scripts. We compare those results as a validation scenario.

#### Scenario #1: conversational exploration of data

The dataset used in this scenario consists of a single 15-minute recording of 4 four mice **(Fig.2)**. It is the original demo dataset^1^ of Live Mouse Tracker (**Fig.2a**). A 30-minute exploratory analysis of the data was performed; and is condensed into an accelerated 5-minute video available here: https://micecraft.org/lmt/mcp/. A website (https://micecraft.org/lmt/lmt-mcp-demo1/) was also generated with the LLM to present the output of the analysis and display details for each figure that you can follow while reading this section. Before using the LLM, the user simply launched the LMT-MCP application, and loaded the database into it. No specific installation of any additional software, apart from the LLM was required. In the following, we provide the prompts submitted to the LLM, and detail the MCP-SIDE MECHANISMS that the LLM uses.

**Figure 2.**
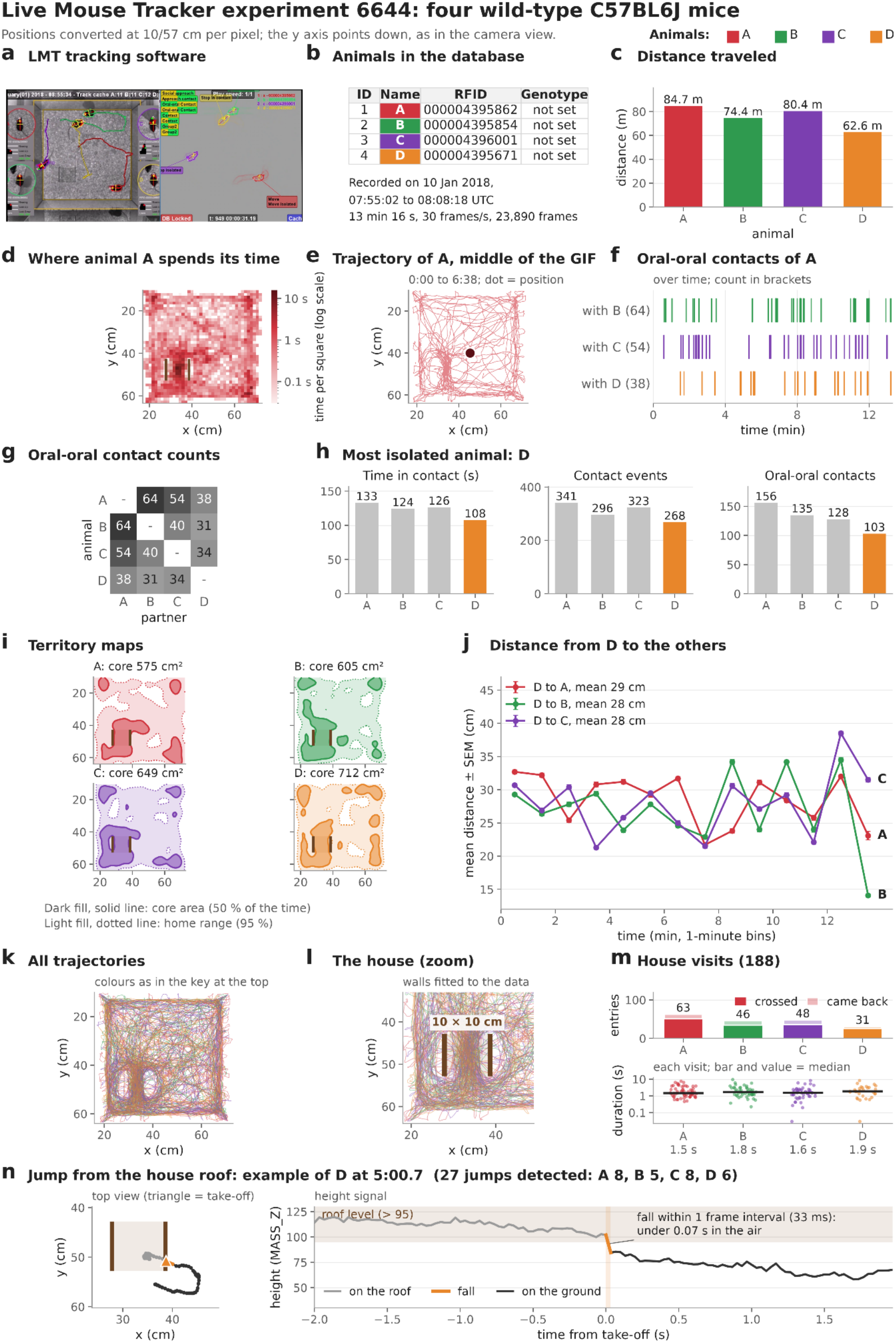
Interactive LLM output using the Live Mouse Tracker experiment. Animal colours: A red, B green, C purple, D orange. (a) LMT tracking & labelling. (b) Animals in the database. (c) Distance traveled. (d) Time spent by A in each part of the cage. (e) Trajectory of A up to the middle of the recording. (f) Oral-oral contacts of A over time. (g) Oral-oral contacts per pair. (h) Social contacts per animal. (i) Territory maps: core area (dark) and home range (light). (j) Distance from D to the other mice over time. (k) All trajectories. (l) The house. (m) House entries and duration of each visit. (n) Example of a jump from the house roof by D: top view and height signal, with the fall in orange.

- *“list the animals in the database”*. Because this is the first prompt, the system has no history, it therefore connects to the MCP automatically to find which database is loaded with the graphical interface of the software using the MCP exposed tool `get_loaded_database`. It retrieves the animals via `list_animals` which returns the lists of the animals with their RFIDs. (**Fig 2b**). Note that one can load additional databases just by prompting *“load database <filename>“*.
- *“duration of the experiment”*. The LLM retrieves the database structure via `get_database_info.` Therefore it uses the FRAME table in the data to provide the exact date and duration of the experiment. Then one of the most common questions asked by biologists concerns the activity and location of the animals, we thus submitted the following questions:
- *“What is the distance traveled by the animals ?*”. The LLM provides the results and the method used for the computation of the distance (**Fig 2c**). *“provide a heatmap of the location of animal A”*. We here specifically refer to one animal only and obtain its heatmap. The LLM uses the MCP-`INSTRUCTIONS` which provides a set of rules read at each session explaining the use of the MCP, the LMT data guidelines and its pitfalls. The INSTRUCTIONS specify that `“The y axis points down (camera view)”` therefore the graph is oriented the same way as the cage seen by the observer (**Fig 2d**).
- *“provide the whole trajectory of the animals A as a GIF of 30s duration”*. The GIF is an image animation format, therefore the LLM provides an animation of the trajectory of the animal A using the DETECTION table (**Fig 2e**). Social interaction events are also important to biologists, we thus submitted the following questions:
- *“give me the oral-oral events of the animal as a timeline”*. The LLM query the MCP with the tool `list_event_types` which provides the list of events available in the database. It can then retrieve the oral-oral events of the animals A with all its other conspecifics and uses the `INSTRUCTIONS` to handle the events table. (**Fig 2f**).
- *“give me the oral-oral event count matrix”* results in **Fig 2g**. The previous questions were driven and reflect immediate access to the data in the database.
- *“Which animal is the most isolated?”*. For this higher level question, the LLM uses the MCP `list_event_types tool` to provide the list of events of LMT, and the LLM uses the built in event within the database to provide an answer based on the time in contact, number of contact events, and number or oral-oral contacts, taking benefits of the `INSTRUCTIONS` directive `“The events can be analyzed in term of number, duration and frequency”` (**Fig 2h**).
- *“create a figure like a territory map for each animal, and one map with all the animals”*. This result is in **Figure 2i**.
- *“what is the distance between animal D and the others ? Plot the graph against time, use bin of*
- *minute”* use the `INSTRUCTIONS` directive “`for the distance between animals, use the mass center X and Y`” (**Fig 2j**).
- *“generate a figure of all the trajectories”* (**Fig 2k**) used the table DETECTION.
- *“the area in the bottom left corner contains a house. It is composed of two walls that the animals cannot cross, and the animals tend to go through quite often. The house is 10×10cm. Draw it on a figure. Fit walls to the data*.*”*. In this prompt, we describe an object , and the LLM finds it at the correct expected location **(Fig 2l)**.
- *“how many times did the mice enter the house?”* provides a count of the number of mice entering the house, number of straight through. For duration: the total time inside the house, the mean visit, median visit and longest one (**Fig 2m**).
- *“The animals can jump from the roof of the house to the ground, find those events and provide a gif as an example of the event”*. With this prompt we asked for a complex event, and we also requested a visual graph to confirm this event (**Fig 2n**).
- *“add this event as “jump from the house” in the database”* used the database information to put the event within the database. Therefore the event can be validated using the LMT dataplayer which shows the events and the tracking for manual validation. Of the 27 detected events, 26 were confirmed. The validation expert judged the remaining one ambiguous.
- *“generate a report with figures as an html page, use plotly for the graphs. Compile the questions we had in this conversation to create sections*.*”*. This provided the website located at https://micecraft.org/lmt/lmt-mcp-demo1/. This query only was executed on high effort.
- *“add all the python scripts to reproduce the data, and the data as* .*xlsx file for each figure”*. For reproducibility purposes, we add the scripts and the data. Note that the scripts are documented and run offline.

In total the LLM evaluated that this scenario consumed 2 kWh, meaning 0.05 to 0.8kg of CO_2_ depending on the nature of the energy used. The LLM wrote python scripts that were launched locally, which limits the LLM energy cost.

#### Scenario #2: full dataset analysis and validation

We used LMT-MCP to reanalyze an LMT experiment that our group had previously analyzed with conventional Python scripts created by an engineer. The paradigm is to explore the social interactions of two 12-week-old female mice with the same genotype in a cage of 50cmX50cm during 2 days (night and daytime). After quality control, the set of experiments included three groups of mice: *Shank3*^*Δ4-22+/+*^ (16 pairs, n = 32 mice), *Shank3*^*Δ4-22*^+/− (12 pairs, n = 24 mice) and *Shank3*^*Δ4-22*^−/− (11 pairs, n = 22 mice). Each pair of mice was recorded for approximately 47 h.

SHANK3 is a scaffolding protein of the postsynaptic density at excitatory synapses. In humans, *SHANK3* haploinsufficiency is a major cause of autism and causes Phelan-McDermid syndrome. This syndrome is characterized by severe speech and language delay, intellectual disability, low muscle tone (hypotonia), global developmental delay and autistic behaviors [17,18]. *Shank3* knockout mice are used as a mouse model of autism [19,20]. The *Shank3*^*Δ4-22*^ Knock-Out (KO) model is a full KO [20] while other *Shank3* KO models have some protein isoforms left.

We loaded 41 LMT SQLite files. LMT-MCP first ran a quality control of the data: it checked that no file was duplicated, identified the genotype of each group and the duration of each experiment. A duplicate database and a 33.6-hour recording were removed for the subsequent analysis.

Our aim was to analyze locomotion and social interactions in each pair of mice and to compare behaviors according to genotype. Our first set of prompts was therefore:

*1- This dataset contains LMT data from three groups of mice corresponding to three genotypes: Shank3+/+ (WT), Shank3+/− (Het) and Shank3−/− (KO). For the first 30 min, the first night, the first day and the second night of the experiment, extract the following data for each genotype: locomotor data (total distance and distance while active, in cm), mean speed (excluding inactivity periods) and inactivity time (s). For distance calculations, remove the data masked by the “flickering” event. For the same periods, extract the number and duration of contacts (total contacts, oral-oral contacts, oral-genital contacts, side-by-side contacts and side-by-side contacts in opposite directions) and of approaches (train2, approach rear, follow zone). Use one value per pair for nose-to-nose and side-by-side contacts. For each parameter, make a graph comparing the genotypes, with the appropriate statistical analysis (and explain the choice of test), in the style of the following graph (an image of a GraphPad Prism graph was inserted here). Write a legend for each graph and show all the code used*.

LMT-MCP detects the night and day periods automatically from clock time. We obtained exactly the results we had requested, organized in a folder. The statistical procedure it chose is as follows: Normality is tested in each group (Shapiro-Wilk) and homogeneity of variances with Levene’s test. If all tests pass, a one-way ANOVA followed by Tukey’s HSD is used. Otherwise, a Kruskal-Wallis test followed by Dunn’s test with Holm correction is used. The statistical unit is the mouse for per-mouse parameters (ignoring that the two mice share a cage) and the pair for per-pair parameters. This represents a classical way for behaviorists to analyze 3-groups data in classical experiments. However, LMT-MCP also points out a limitation: “the unit of analysis is the animal, but the two mice in a cage share their environment and are not truly independent. A stricter version would average each cage (n = 16, 12 and 11 cages) or use a mixed model with “cage” as a random factor.” We therefore ask for the latter option and obtain the mixed-model analysis after a re-run.

We then asked for a website that brings together all the results generated so far. We submitted the following set of prompts:

*2- With high effort, create a static website in English, in the style of a report, with several tabs organized by theme. Make the graphs with Plotly and include all the analyses done previously. Run the statistics with a mixed model. Add some debugging to the pages: a button called “Additional information” that expands extra text, so I can check that the counts are correct. For the distance calculations, remove the data masked by the “flickering” event. For each criterion, add an indicator of each group’s level compared with the others. Provide the xlsx file and the code for each figure, with a simple download button. Add a radar plot underneath to compare the groups. Also create general top-level sections that group the social criteria and the motor criteria. Put the experiment summary at the top of the page, with the databases, cages and mice, in an “Experiment info” tab. Add a “Main message” tab with a summary text and the graphs (with xlsx and code) of the data that differ significantly. This tab must be shown by default when the site opens. “In the “Main message” tab, add two PheWAS figures side by side at the top, using only the “whole experiment” data: on the left, the Shank3+/+ vs Shank3−/− comparisons, and on the right, the Shank3+/+ vs Shank3+/− comparisons. Put the different measured parameters on the x-axis, with a different color for each parameter, a clear separation between social and motor parameters, and the behaviors labeled along the x-axis. Put −log10(p-value) on the y-axis. Use the style of this graph (here the user drags an example of a PheWAS in the console). Below them, make two PheWAS plots in the same style with all the data except “whole experiment”, with Shank3+/+ vs Shank3−/− placed above Shank3+/+ vs Shank3+/−. “here the user provides a file* .*xls containing the analysis of the same LMT experiments made by an engineer with Python coding”: this file contains an analysis of the same data done by an engineer in Python. “d1” and “d2” correspond to the first 24 h and the second 24 h, and “all” corresponds to the whole experiment. Make the graphs from these data, checking that they are the same animals as in your analysis. Also analyze your own data over the first 24 h and the second 24 h. Then add a “Validation” tab to the website, with mirrored graphs comparing the engineer’s analyses and your analyses for each common variable for 0-24h, 24-48h and the whole experiment. Mark each comparison with a green dot if the two are similar and an orange dot if they differ (based on the correlation). Use the same units for both analyses. Put a summary table at the top of the page, and make each table cell clickable so that it takes you to the corresponding graph. Add this link as a button in the top-right corner of the website: https://micecraft.org/lmt/mcp/ Add a banner at the very top of the website that stay fixed when scrolling, like the one on the site hosted at https://micecraft.org/lmt/lmt-mcp-demo1/, containing “Live Mouse Tracker MCP report generation demo” and a “Go to LMT-MCP” button. Add to the subtitle that the results were compared, for validation, with an analysis done by a human analyst*.*In the main menu, highlight the “Validation” button. Add a “Prompts” tab with a compact version of the prompts received to generate the analyses and the website. Add a figure that summarizes the essentials of the main message, to the right of the main message, with clickable cells linked to the detailed figures*.*”*

The resulting website is available here: https://micecraft.org/lmt/lmt-mcp-demo2/#general. The LMT-MCP analysis suggests that **social behavior** is atypical in female *Shank3*^*Δ4-22*^−/− mice (fSh3-/-) compared with their wild-type female littermate. The most prominent difference was in oral-oral contact: over the whole experiment, fSh3-/-pairs spent more time in oral contact than fSh3+/+ (p = 0.0013) and fSh3+/- (p = 0.0016) pairs. We then make a closer look at the first 30 min, a period characterized by novelty and stress. We observed that fSh3-/-make more close contacts than fSh3+/+ pairs (longer oral-genital contacts, more and longer side-by-side contacts). Over longer periods the differences narrowed to two behaviors: the excess oral–oral contact persisted during both nights and across the whole experiment (vs. both fSh3+/+ and fSh3+/− pairs), and fSh3−/− mice followed their cagemate more often (FollowZone) during the first 30 min, during both nights and across the whole experiment, but not during the day. The only difference that stays significant after correction for multiple analysis is the increased time in oral-oral contacts in fSh3-/-mice. This increase in oral-oral contact time was observed in other models of Shank3-deficiency in LMT experiments [21,22,23]. fSh3+/-mice do not differ from fSh3+/+ mice on any social criterion. The **motor activity** is slightly reduced in fSh3+/-mice, but does not reach statistical significance after correction for multiple testing. fSh3+/-mice travel less distance than fSh3+/+ and fSh3-/-mice during both nights and over the whole experiment. They are active for less time than fSh3+/+ mice in night 1 and than fSh3-/-mice in both nights and over the whole experiment. During the first day they spend more time inactive than fSh3+/+ mice. fSh3-/-mice are not hypoactive in this experiment.

The “validation” tab shows 100% similarity between the engineer analysis and the LLM analysis thus validating the LMT-MCP for this in depth analysis of a large dataset in a short time and without prior coding knowledge.

The results are summarized in **Figure 3**.

**Figure 3.**
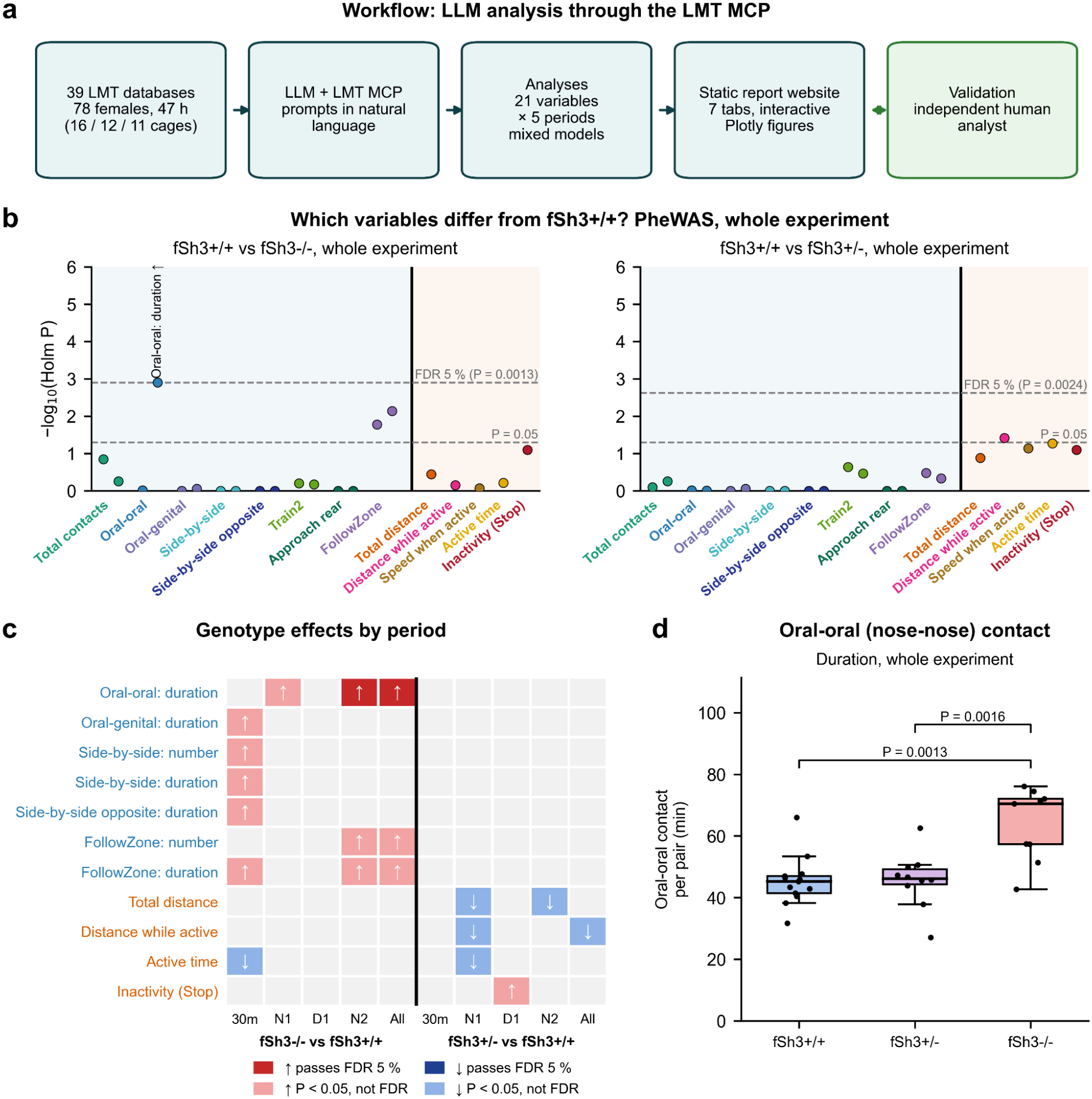
LLM-generated analysis and report of Live Mouse Tracker recordings of female Shank3-mutant mice (scenario #2) (a) Workflow of scenario #2: full dataset analysis with LMT-MCP and validation. (b) PheWAS-style plots for the whole experiment: fSh3+/+ vs fSh3-/-(left) and fSh3+/+ vs fSh3+/-(right). Each point is one variable, grouped and coloured by behavior (number (left) and duration (right) of the same behavior share a colour. Dashed lines: P = 0.05 and the Benjamini-Hochberg false discovery rate threshold (FDR 5 %), computed over the 21 variables of each plot. Labelled point: variable above the FDR threshold (↑, higher in the mutant). Only oral-oral contact duration in fSh3-/-mice passes the correction for multiple testing (c) Summary of genotype effects across periods. Each row is a variable with at least one significant difference from fSh3+/+ (social variables in blue, motor variables in orange); columns are periods (30m: first 30 min; N1: night 1; D1: day 1; N2: night 2; All: whole experiment), for fSh3-/-(left) and fSh3+/-(right) versus fSh3+/+. Coloured cells: Dark red ↑ / dark blue ↓: higher / lower than fSh3+/+ and passing the Benjamini-Hochberg FDR 5 % threshold of the corresponding PheWAS plot. Pink ↑ / light blue ↓: Holm-corrected P < 0.05 but above the FDR threshold. Grey: P ≥ 0.05.. (d) Oral-oral contact duration over the whole experiment, per pair (one value per cage; fSh3+/+ n = 16, fSh3+/-n = 12, fSh3-/-n = 11 cages, minus cages excluded for incomplete event computation). Boxes: median and interquartile range. P values: Holm-corrected pairwise contrasts of a linear model on log(x + 1).

The one-day analysis of scenario #1 used about 10M tokens. That cost roughly €7 and emitted about 130 g of CO_2_. Over a year of 250 working days, this comes to about €1,750 and 32 kg of CO_2_ per researcher. That matches the Celsius Project’s estimate of 10–50 kg CO_2_eq per user of agentic AI [24]. This footprint probably reflects higher productivity, but users should still keep it in mind. To help them estimate it, the `/mcp_ _lmt-eye ecological_impact` command gives a rough, formatted estimate of an analysis’s impact.

## Conclusion & Perspectives

As expected from the scientists and editors, AI is supposed to play an increasing role in science [25,26]. This still raises some new questions in terms of methodology [14-15] but may open a new opportunity for dialoguing with science [13], understanding [16] or reproducibility [7]. This new tool makes LMT data analysis accessible for experimentalists, which was clearly needed by the teams equipped with LMT.

However, it also raises new questions for the community that are tied to the use of AI. Some are practical such as data ownership, security or local AI model deployment. Others are scientific, such as labor division among the teams or ensuring that the scientific critical oversight is kept on the analysis. Still others go deeper, such as the ecological impact. Those questions are now opened to a larger community, and the channel is open on the LMT discord channel: https://discord.gg/zWDHNf9eHM.

## Author’s contributions

G.N. programmed the MPC, X.M. coded the LMT-Eye application, N.L., S.C. took care of the animals, M.M. and B.F. performed LMT experiments and autonomous test analysis, T.B. and F.d.C. provided constraints and original ideas. G.N., F.d.C., B.F. & T.B. wrote the article.

## Acknowledgements

This work was funded by Institut Pasteur, Université Paris Cité, the CNRS. This work received support from the French government, managed by the National Research Agency (Agence Nationale de la Recherche), through the Inception program (Investissement d’Avenir grant ANR-16-CONV-0005), under the France 2030 program (ANR-23-IAHU-0010), and the AutismSHANK project (ANR-23-CE16-0013). This work was also funded by the DIM C-BRAINS initiative.

## Supplementary materials

### Integrated Instructions

The `INSTRUCTIONS` is the pre prompt provided to the LLM by the MCP. The LLM will use it when connecting to the MCP. This pre prompt will change along the MCP versions.

~~~
Access to Live Mouse Tracker (LMT) experiments opened in LMT-EYE. Without ‘database_path’, tools use the database currently loaded in LMT-EYE. All tools are read-only.
~~~

#### LMT database conventions

~~~
•Positions (MASS_X/Y, FRONT_*, BACK_*) are in pixels. Convert with 10/57 cm per pixel for mice and 20/57 for rats. The y axis points down (camera view).
•Frames are usually at 30 frames/s (see get_database_info). Time 0 is the first frame of the FRAME table.
•MASS_Z is a relative height of the animal from the ground in millimeters: low on the floor, higher on raised objects such as a house roof. Check the levels in each setup.
•Pairwise events (e.g. ‘Oral-oral Contact’, ‘Contact’) are stored once per animal of the pair (A->B and B->A). Merge them before counting contacts between two animals.
•A centre-of-mass step longer than 85.5 px between two consecutive frames is a tracking error. Exclude it from distances.
•Even when an animal is stopped, the tracking may still move a little around the animal, which provides overestimated distances.
•Identity switches can happen when two mice are close: the tracked position jumps by several cm in one frame. Check for them before trusting a rare event.
•For data analysis report, prefer a webpage using plotly for figures.
•If events are missing in the database, the rebuild all events function compute them. The table
LOG displays in each row the last time the event has been computed, and for which time range (in
frame).
•In the table events, the IDANIMAL* represents the animals in the order of the event description (e.g. Oral-genital Contact means contact of the oral part of IDANIMALA with genital of
IDANIMALB). One exception for nest3*: the idanimalA is the one that is not in the nest. Nest3 only work if 4 animals are in the database.
~~~

#### Quality control

~~~
•The frame should be at 30fps
•Check duplicated databases
~~~

#### Analysis

~~~
•The events can be analyzed in term of number, duration and frequency
•For the distance between animals, use the mass center X and Y
•For the sake of reproducibility, generate python codes
•It is not parameters but variables
•Stats: mice of a same experiment are not independent, the experiment is the statistical unit
~~~

#### Color conventions

~~~
•Animal A is red, B green, C purple, D orange
~~~

### Integrated prompts

The integrated prompts are prompts accessible via the “/” command in LLMs. When one type /lmt in the prompt, the user can see the whole set of functions we provide.

#### Ecological impact

This prompt is called with the */lmt-eye_ecological_impact*:

~~~
Using the current conversation with the user, estimate the impact in kWh of the usage, and the volume of water consumption. Display the difference between the energy used on the server side, and on the user’s computer. Provide good practices to save energy.
~~~

#### Integrated instructions

This prompt is called with the */lmt-eye_integrated_instructions_list*:

~~~
Display the instructions of this MCP to the user.
~~~

## Footnotes

1 https://micecraft.org/lmt/download/20180110_validation_4_ind_Experiment_6644_e.sqlite

## Notes

https://micecraft.org/lmt/mcp/

### Competing Interest Statement

The authors have declared no competing interest.

